# Characterization of the gut microbiota in the Anjak (*Schizocypris altidorsalis*), a native fish species of Iran

**DOI:** 10.1101/735241

**Authors:** Mahdi Ghanbari, Mansooreh Jami, Hadi Shahraki, Konrad J. Domig

## Abstract

The antiquity, diversity and dietary variations among fish species present an exceptional opportunity for understanding the variety and nature of symbioses in vertebrate gut microbial communities. In this study for the first time the composition of the gut bacterial communities in the anjak was surveyed by 16S rRNA gene 454-pyrosequencing. Anjak (*Schizocypris altidorsalis*), is a native benthopelagic and commercially important fish species in Hamun Lake and Hirmand River system in Sistan basin, Iran. Thirteen 16S rRNA PCR libraries were generated, representing 13 samples of Anjak gut microbiota. After quality filtering, a total of 137 978 read pairs remained from the V3–V6 16S rRNA gene regions, each measuring 680 bp, and were assigned to bacterial OTU’s at a minimum sequence homology of 97%, resulted in total 608 OTUs. The results provided evidence of the presence of a very diverse microbial community present in this fish. The majority of sequences belonged to members of the Firmicutes; however, members of the Proteobacteria, Tenericutes, Bacteroidetes and Cyanobacteria were also well represented in the anjak gut microbiota. The most abundant classes recorded in the samples were uncultured CK-1C4-19 class of Tenericutes, Flavobacteriia, α-Proteobacteria, Erysipelotrichi, Synechococcophycideae, γ-Proteobacteria, and Clostridia. The presence of many different types of bacteria in the anjak is not unexpected since it has been shown that eukaryotes with a detritivorous and planktivorous diet have a higher microbial diversity compared to omnivorous and herbivorous fish species. Many of these microbes might be of high physiological relevance for snow trout and may play key roles in the functioning of a healthy gut in this fish.

## Introduction

Indigenous microbiota play a crucial role in their vertebrate hosts. In fact, it is increasingly clear that the intestine is a multifunctional organ system involved in the digestion, resorption and absorption of food components, in electrolyte balance, endocrine metabolic regulation of and immunity against pathogens (Austin 2002; Ghanbari et al. 2015; Ghanbari et al. 2009; Ghanbari et al. 2017; Jami et al. 2015; Nayak 2010). Comprising a large diversity of approximately 28,000 species, fish include nearly half of all vertebrate species and exhibit a wide variety of physiologies, ecologies and natural histories (Wong and Rawls 2012). The antiquity, diversity and dietary variations among fish species present an exceptional opportunity for understanding the variety and nature of symbioses in vertebrate gut microbial communities. Studies have shown that the composition of the gut microbiota varies among fish species, with the predominance of Proteobacteria and Firmicutes, but the bacterial composition may vary with age, individuals, developmental stage, nutritional status, environmental conditions, and the complexity of the fish digestive system (Ghanbari et al. 2015; Ghanbari et al. 2017; Wu et al. 2013).

Next generation sequencing (NGS) platforms offer the possibility to explore the diversity of microbial communities on an unprecedented scale, and indeed have been principally used to study the microbial ecology of a wide range of environments including the gut microbiota (Ghanbari et al. 2015; Kraler et al. 2016). Numbers of bacterial 16S rRNA sequences derived from a sample can be assigned to individual genera/species thus giving a semi-quantitative estimation of the relative abundance of each microorganism present (Ghanbari et al. 2015). NGS platforms such as 454-GS FLX can generate millions of paired-end microbial 16S rRNA gene sequences from a single sample, offering an extremely detailed picture of the structure of the gut microbial communities and allowing researchers to test more complex hypotheses concerning the microbial communities that abound in gut ecosystems than lower resolution molecular and culture-based methods (Ghanbari et al. 2015).

Anjak (*Schizocypris altidorsalis*), is a native benthopelagic and commercially important fish species in Hamun Lake and Hirmand River system in Sistan basin, Iran (Coad and Ville 2005; Jami et al. 2015). Together with snow trout (*Schizothorax zarudnyi*), anjak make up the most favourite source of animal protein for Sistan residents. It belongs to the family *Cyprinidae* and sub-family Schizothoracinae, which are widely distributed in the Himalayan and sub-Himalayan region and the rest of Asia (Jami et al. 2015). Almost nothing is known about the nature and biodiversity of intestinal microbiota in this fish. Hence in this study the composition of the gut bacterial communities in the anjak was surveyed by 16S rRNA gene 454-pyrosequencing.

## Materials and methods

### Sample collection and preparation

For this study, fish samples representing anjak were obtained from Chahnimeh reservoirs, Sistan, Iran (61° 63⍰ - 61° 73 ⍰longitude and 30° 74⍰-30° 84⍰ latitude) by fishing. Thirteen adult fish with an average weight of approx. 200 g were sacrificed using high doses of anaesthesia benzocaine (Sigma Aldrich, UK). All instruments, surfaces, and the exterior of each fish were treated with 70% EtOH and instruments were flame-sterilized prior to dissection. After opening the body cavity, the intestines were separated, dissected, and were placed directly into sterile 2 mL screw cap tubes containing sterile containing phosphate-buffered saline (PBS)/glycerol (50:50, v/v) and stored at -20°C.

### DNA extraction

Metagenomic DNA was extracted from frozen samples using the RBB+C method (Yu and Morrison 2004) with some modification. Briefly, 0.25 g of gut samples were resuspended into 710 μL of 200 mM NaCl, 200 mM Tris, 20 mM EDTA plus 6% SDS. After the addition of 0.5 mL of 0.1-mm zirconia beads (BioSpec Products) cells were lysed by mechanical disruption with a TissueLyser LT (Qiagen) for 5 min. Then, the tubes were incubated at 70°C for 15 min. Samples were centrifuged at 4 °C for 3 min at 16 000 × g, and the aqueous phase was collected and DNA in the aqueous phase was precipitated by the addition of an equal volume of isopropanol and 0.1 volumes of 3 M sodium acetate, pH 5.5 (Carl Roth). After overnight extraction at −20 °C, samples were centrifuged for 20 min at 4 °C at 16 000 × g, and the supernatant was removed. After washing once with 0.5 mL of 100% ethanol, the pelleted DNA was allowed to dry and was re-suspended in TE buffer and treated with RNAse and Proteinase K. It was cleaned with QIAGEN buffers AW1 and AW2 and eluted in 50 μL of AE buffer (QIAamp DNA Stool Mini Kit, QIAGEN, UK). DNA was visualized on a 0.8 % agarose gel and quantified using the Qubit 2.0 fluorometer (Invitrogen, Carlsbad, CA, USA). Extracted DNA was stored at −80 °C.

### Molecular analysis

Approximately 10⍰ng of DNA from each of the obtained 13 mgDNA samples was used to build each targeted library. Samples, negative extraction controls, and reagent blanks were PCR-amplified using primer constructs containing the universal 16S rRNA V3-V6 region 338F/1061R primers (Ong et al. 2013), tagged with the Lib.A and Lib.B Roche 454 Titanium sequencing adapters. Each PCR reaction contained 14.75⍰μL molecular grade water, 5⍰μL 5x KAPA HiFi fidelity buffer, 0.75⍰μL 10 mM KAPA dNTP Mix, 0.75⍰μL 10⍰μM primer 338F, 0.75⍰μL 10⍰μM primer 1061 R, 0. 5⍰μL KAPA HiFi hotstart DNA polymerase (1⍰U/μL) and 2.5⍰μL of DNA template for a total volume of 25⍰μL. The temperature profile for the reactions included an initial denaturation step at 95°C for 3 min, followed by 26 cycles of denaturation at 98°C for 20 s, annealing at 66°C for 15 s and extension at 72°C for 20 s. The final extension step was performed at 72°C for 60 s. The amplified PCR products were visualised by 2% low-melting gel electrophoresis. For each sample, the PCR products of two successful amplifications were pooled and purified with PureLink PCR purification Kit (Invitrogen, CA, USA). A Qubit 2.0 fluorometer (Invitrogen, Carlsbad, CA, USA) was used to quantify the resulting pooled DNA. Equimolar amount of the PCR products from each sample were pooled together and sequenced unidirectional on a 454 GS FLX Titanium platform applying the so-called GS FLX+ system (Roche Applied Science, Indianapolis, IN, USA) at the Eurofins-Genomics centre (Ebersberg, Germany). The PCRs from negative controls yielded negligible DNA concentrations during Qubit 2.0 quantitation, indicating contamination was not a problem.

### Bioinformatics

Sequence de-multiplexing and bioinformatic processing of the 454 dataset was performed using aspects of the UPARSE (Edgar 2013) and QIIME v 1.9.1 (Caporaso et al. 2010) pipelines. Initial quality filtering of 454 sequences excluded all sequences <200 or >1000 bp, with any primer mismatches, with a homopolymer run 10 bp. A de novo database of ≥97% similar sequence clusters was created in USEARCH (Version 8); (Edgar 2013) by (i) quality filtering sequences using a “maxee” value of 1 (i.e., sequences with a predicted error rate of 1 bases per sequence were discarded) and “fastq_trunclen 680”, (ii) dereplicating identical sequences, (iii) removing singleton sequences, and (iv) clustering those sequences (Edgar 2013). Chimeras were checked against the ‘‘Gold’’ database based on UCHIME (Edgar et al. 2011). Raw demultiplexed sequences were then mapped against these de novo databases to generate counts of sequences matching clusters (i.e., phylotypes) for each sample. Taxonomy was assigned to each phylotype by using the RDP algorithm with a confidence threshold of 0.8 and trained on the Greengenes database (DeSantis et al., 2006), updated May, 2013. The raw sequence data are available in the Sequence Read Archive at the National Center for Biotechnology Information (BioProject accession no. PRJNA335697).

## Results

### Sequence statistic

Thirteen 16S rRNA PCR libraries were generated, representing 13 samples of anjak gut microbiota. After quality filtering, a total of 137 978 read pairs remained from the V3–V6 16S rRNA gene regions in this study, each measuring 680 bp, and were assigned to bacterial OTU’s at a minimum sequence homology of 97%, resulted in total 608 OTUs.

### Microbial community composition of snow trout intestinal lumen

Figure 1 illustrates the abundance of major (>1%) phyla and classes observed in the gut samples. Details of microbial taxon composition, diversity, and frequencies were given in supplementary data 1. Bacteria representing 20 phyla were found in the anjak gut samples. The majority of sequences belonged to members of the Firmicutes (mean sequence abundance 26.3%); however, members of the Proteobacteria (19.6%), Tenericutes (13.9%), Bacteroidetes (12.0%) and Cyanobacteria (10.8%) were also well represented in the gut microbiota. Other phyla including Actinobacteria (4.9%), Fusobacteria (3.3%) and Chlamydiae (1.3%) were relatively less pronounced (Fig. 1a), with the rest of phyla were present in low abundance (<1%). The most abundant classes recorded in the samples, in terms of mean sequence abundance, were uncultured CK-1C4-19 class of Tenericutes (13,9%), Flavobacteriia (11.9%), α-Proteobacteria (10,2%), Erysipelotrichi (8.2%), Synechococcophycideae (7.1%), γ-Proteobacteria (7.1%), and Clostridia (6.6%) with the rest of classes were present in lower sequence abundance (<5%, fig. 2b, supplementary data 1).

**Fig. 1.**
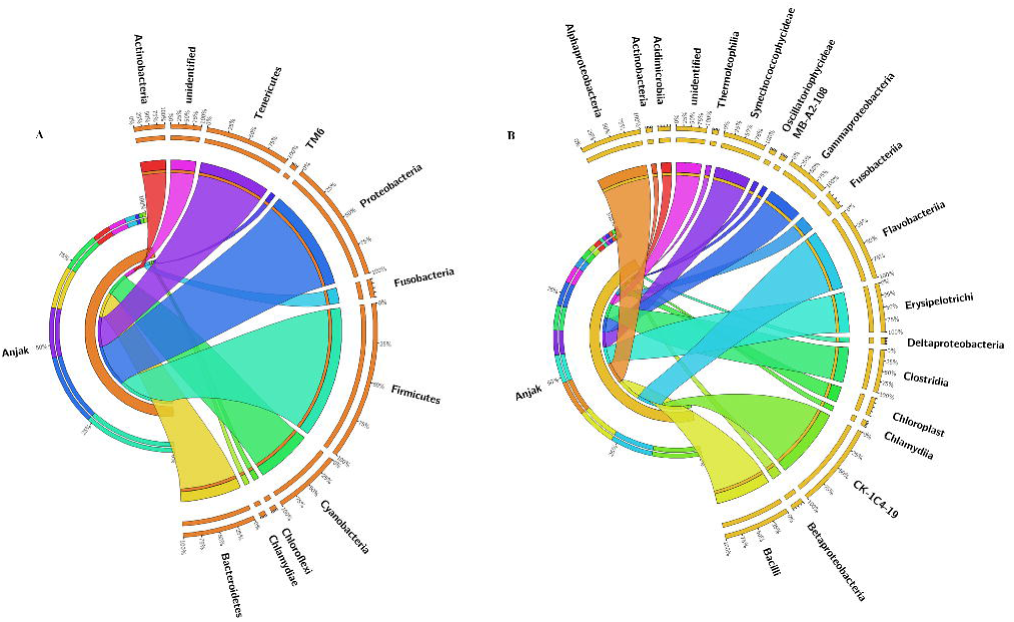
Circular representation of microbial communities in anjak at phylum (a) and class (b) level. Phyla and classes with relative abundance lower than 1% were not shown

A large proportion (60%) of the reads in all the libraries could not be classified at the genus level. The most abundant genera, based on an abundance cut-off of 1%, in the anjak gut microbiota were sequences related to CK-1C4-19_group (Tenericutes, CK-1C4-19, 13.9%), *Flavobacterium* (11.7%), *Holdemania* (8.2%), *Synechococcus* (7.1%), unclassified bacterial group (5.2%), *Lactobacillus* (5.1%), *Clostridium*_XI (5.1%), unclassified Rhizobiales (3.9%), *Pediococcous* (3.5%), *Aeromonas* (3.2%), uncultured u114 (Fusobacteria, Fusobacteriia, Fusobacteriales, Fusobacteriaceae, 2.5%), unclassified SC-I-84 group (Proteobacteria, Betaproteobacteria, SC-I-84, 1.4%), unclassified Legionellaceae (1.4%), *Rhodobacter* (1.3%), unclassified C11 group (Actinobacteria, Acidimicrobiia, Acidimicrobiales, C11, 1.2%), *Pleomorphomonas* (1.1%) and unclassified Stramenopiles (1%). Gram-positive bacteria (*Bacillus, Carnobacterium, Enterococcus, Lactobacillus, Lactococcus, Micrococcus, Pediococcus, Streptococcus*, and *Weissella*), and Gram-negative bacteria (*Aeromonas, Alteromonas, Photorhodobacterium, Pseudomonas*, and *Vibrio*) have been used successfully as probiotics in aquaculture (Balcázar et al. 2006). In the present study, reads related to the genera *Bacillus, Lactobacillus, Lactococcus, Micrococcus, Streptococcus, Pediococcus, Aeromonas, Pseudomonas*, and *Vibrio* were detected in the gut microbiota of anjak, although the reads were in relatively low abundance (<6%).

## Discussion

It is widely acknowledged that the vertebrate gut microbial communities play critical roles in host immune system and digestive system (Ghanbari et al. 2015; Kraler et al. 2016; Liu et al. 2016), which is now attracting increasing attention in the fish research. Fish gut-associated microbial communities exert some pronounced effects on host nutrition, physiology, and growth (Ghanbari et al. 2015; Ye et al. 2014). Although studies on gut microbiota in some carp species have been reported (Han et al. 2010; Jami et al. 2015; Li et al. 2015a; Li et al. 2015b; Li et al. 2013; Li et al. 2014; Liu et al. 2016; Ni et al. 2014; Ray et al. 2009; Schaus et al. 1997; van Kessel et al. 2011; Wu et al. 2012; Wu et al. 2013), no knowledge exists about gut microbial community in anjak. Previous studies have shown that 454-pyrosequencing has a higher capacity than culture-dependent methods to explore bacterial richness, indicating that pyrosequencing is an efficient way to access microbial diversity (Ghanbari et al. 2015). This study demonstrated, to the author’s knowledge, the first successful application of pyrosequencing using the 454 GS FLX+ to investigate the gut microbial community in an Iran’s native fish species with economical value.

Surveys on the vertebrate gut samples, especially terrestrial mammalian gut microbiota, suggest that Firmicutes and Bacteroidetes are numerically the most dominant phyla (Kraler et al. 2016; Ley et al. 2008; Qin et al. 2010; Wu et al. 2012). The present study suggests that the anjak gut microbial community is dominated by Firmicutes, followed by Proteobacteria, Tenericutes and Bacteroidetes and Cyanobacteria. The results are generally consistent with those of Han et al. (2010), Wu et al. (2010), Roeselers et al. (2011); Wu et al. (2012), Tetlock et al. (2012),, Lyons et al. (2015), Liu et al. (2016), Zac Stephens et al. (2016) and Ghanbari et al. (2017), who demonstrated that species from the phyla Proteobacteria, Firmicutes, Bacteroidetes, and Tenericutes are commonly among the most dominant members present in the GI tracts of different fish species (For a comprehensive review see (Ghanbari et al. 2015; Llewellyn et al. 2014; Nayak 2010). In fact the presence of the dominating soil phylum such as Proteobacteria, Bacteroidetes and Chloroflexi in the aquatic environment is very common (van Kessel et al. 2011; Wu et al. 2012a; Wu et al. 2013; Ye et al. 2014; Li et al. 2015a).

Within the Firmicutes phylum, Bacilli, Erysipelotrichi and Clostridia were present in the microbiota of the anjak gut (Fig. 1b). Representative genera of these abundant classes, including *Clostridium, Bacillus, Lactobacillus, Pediococcus, Streptococcus* and *Staphylococcus* spp., have been frequently reported in the microbiota of various fish such as carps (Liu et al. 2016; Rawls et al. 2006; Ray et al. 2010; van Kessel et al. 2011; Wu et al. 2012; Wu et al. 2013) and rainbow trout (Lyons et al. 2015; Wong et al. 2013). Previous studies on the functional characteristics of the members of the Clostridia and Bacilli isolated from gut microbiota have shown that these bacteria posse a wide-ranging metabolic capabilities which able them to utilize various sugars as sole carbon sources (Liu et al. 2016; Ray et al. 2010; Wu et al. 2012). In addition, the majority of the genera belong to these classes in the current study, including *Lactobacillus, Clostridium, Pediococcous, Bacillus* are known to produce useful end-products, (Ghanbari et al. 2013b; Ghanbari et al. 2009; Montoya et al. 2001) which is important in the niche environment being studied here. An interesting feature, given that these genera are examples of the dominant classes present in a single micro-environment, is the fact that one group is aerobic (*Bacillus*), one group is microaerophilic (*Lactobacillus*) and the other is anaerobic (*Clostridium*). This is an ideal situation for a microbial community as the component genera are complimentary rather than competitive in their gaseous requirements. The ability of Bacilli class members like *Bacillus* and *Lactobacillus* sp. to form an extracellular matrix (e.g. polysaccharides) is also beneficial to the microbial community (Ruas-Madiedo et al. 2007; Sims et al. 2011; Ullrich 2009). Given the ability of *Clostridium* species to break down polysaccharides, it is interesting to note the potential dynamic – polysaccharide production and degradation occurring within the same community. *Holdemania* was the most abundant genus in the Firmicutes phylum. The genus is related to Erysipelothrix according to 16S rRNA gene analysis and was shown to be enriched on the chicken (Gong et al. 2002) and pig cecal mucosa (Buzoianu et al. 2012; Gong et al. 2002; Kim et al. 2011; Looft et al. 2014). Although the role of this genus in the intestine is not fully understood, the high presence of the *Holdemania* spp. in gut microbiota has been linked to antibiotic treatments and antibiotic resistance features (Looft et al. 2014; Willems et al. 1997). Willems et al. (1997) reported Holdemania spp. isolated from human gut microbiota have a unique murein type as part of their cell wall structure, which may contribute to insensitivity to some antibiotics.

Within the phylum Proteobacteria, the α-Proteobacteria was the dominant group followed by the γ-Proteobacteria. The class of α-Proteobacteria harbors a miscellaneous set of metabolisms, cellular phenotypes and a wide range of habitats from the ocean floor volcanic environments, to soil, in which they may interact with plant roots, to surface waters of oceans (Pini et al. 2011). The wide variety of genera of this class were found in the current study including phototrophic genera (*Rhodobacter*), symbionts of plants (*Rhizobium*), nitrogen-fixing genera (*Pleomorphomonas*), animal and plant pathogens (*Rickettsia, Agrobacterium*) and also genera able to metabolize C1 compounds (*Methylobacterium, Methylosinus*) (Narrowe et al. 2015; Pini et al. 2011). Beside the unclassified genus from Rhizobiales which is likely originating from plant material (Pini et al. 2011), the genus Rhodobacter had the second highest abundant in the anjak gut. This genus is widely distributed in fresh water as well as marine and hypersaline habitats (Giatsis et al. 2015). *Rhodobacter* species have been reported in the gut of grass carp (Wu et al. 2013), rainbow trout (Mansfield et al. 2010; Wong et al. 2013), Siberian sturgeon (Geraylou et al. 2013), catfish (Di Maiuta et al. 2013) and Atlantic cod (Fjellheim et al. 2012), as well as in the water of aquaculture systems (Petersen et al. 2013). The γ-Proteobacteria class includes a wide range of pathogens, such as *Aeromonas, Aquicella, Legionella, Pseudomonas* sp., *Escherichia coli, Salmonella* sp., *Vibrio* sp., etc (Austin and Austin 2012). However, the majority of species in this class are free-living, and also include a number of nitrogen fixing bacteria and enzyme-producing (amylase, cellulase and protease) bacteria (Austin and Austin 2012; Sharmin et al. 2013), which may explain their high abundance in gut microbiota of in gut microbiota of detritivorous anjak. Most γ-Proteobacteria found in anjak belonged to the *Aeromonas* group (Supplement data 1). Members of this genus are mainly distributed in freshwater and sewage, often in association with aquatic animals. They can cause a diverse spectrum of diseases in both warm- and cold-blooded animals but they also appear to be simply commensal and act as naturally occurring members of bacterial communities in aquatic envrionments including in fish intestines (Ghanbari et al. 2017; Larsen et al. 2014; Ray et al. 2012; van Kessel et al. 2011; Wu et al. 2012).

Within the anjak gut bacterial communities, we identified a highly abundant taxon, CK-1C4-19, that is classified as either a member of the phylum Tenericutes in Greengenes database (DeSantis et al. 2006) or a candidate division at the phylum level according to the SILVA ribosomal RNA gene database project (Quast et al. 2013). OTUs from this taxon have been previously identified as members of the zebrafish (Cyprinidae, *Danio rerio*) (Roeselers et al. 2011; Zac Stephens et al. 2016) and fathead minnow (*Cyprinidae, Pimephales promelas*) gut microbiome (Narrowe et al. 2015) and a relatively small number of sequences exist in 16S rDNA databases, with annotated environmental and host-associated habitats including other cyprinid fishes, ants, lobster, catfish, sediments, and anaerobic digesters.

Another well represented group within the obtained sequences were the Bacteroidetes (12% of obtained sequences), a phylum known for a fermentative metabolism and degradation of oligosaccharides derived from plant material (van Kessel et al. 2011). Almost all the sequences in this phylum belonged to the genus *Flavobacterium* (Fig. 1, 11.7%). *Flavobacterium* members reside in diverse habitats, including freshwater streams, lakes, and sediments, deep wells, glaciers and arctic ice, plants and plant material, soils, freshwater shrimp ponds, marine sediments, seawater, and on marine (Bernardet and Bowman 2006; Loch and Faisal 2015). By far and away, the most plentiful reports of *Flavobacterium* spp. in associated with animals are those that document their presence on fish. For example, *Flavobacterium* spp. have been detected within the intestines (Hu et al. 2007; Huber et al. 2004; Ray et al. 2012), and from internal organs of cool, cold, and warm-water fishes (reviewed in (Bernardet and Bowman 2006; Ray et al. 2012; Starliper 2011). Although some *Flavobacterium* spp. are commensal a number are major pathogens of freshwater and marine fishes (Loch and Faisal 2015). It has been suggested that fish gut isolated F*lavobacterium* sp might be possible contributors to the digestive processes of fish by producing enzymes such as amylase (Skrodenytė-Arbačiauskienė 2000), chitinase (Ray et al. 2012) and protease (Morita et al. 1998; Skrodenytė-Arbačiauskienė 2000).

The high abundance of Cyanobacteria observed likely support their importance as food sources for anjak. In fact Cyanobacteria are known to be important food sources for detritivorous and planktivorous fish (Beveridge et al. 1993; Schaus et al. 1997; Ye et al. 2014). Within this phylum, *Synechococcus* was the most dominant genus observed in the gut microbiota of anjak. The genus *Synechococcus* encompasses cyanobacteria in both freshwater and marine environments and seems to be more abundant in nutrient rich aquatic environments than oligotrophic environments that contain a minimal amount of nutrients (Gao et al. 2014; Whitton 2012).

Actinobacteria, Fusobacteria and Chlamydiae were three low abundant phyla detected within the anjak gut microbiota. Actinobacteria are well known for production of secondary metabolites, of which many are potent antibiotics (Jami et al. 2015). They are widely distributed in both terrestrial and aquatic (including marine) ecosystems, especially in soil, where they play a crucial role in the recycling of refractory biomaterials through decomposition and humus formation (Ventura et al. 2007). Previous studies have reported that the phylum generally makes up a small proportion of bacterial sequences retrieved from fish intestine (Lyons et al. 2015; van Kessel et al. 2011; Wong et al. 2013; Xia et al. 2014; Ye et al. 2014) and may contribute to cellulose degradation in herbivorous fish (Liu et al. 2016; Ray et al. 2012). The Fusobacteria are anaerobic, Gram-negative bacilli that produce butyrate, a short chain fatty acid that is often the end-product of the fermentation of carbohydrates including those found in mucins (Larsen et al. 2014). The presence of these obligate anaerobes bacteria suggest that they may play an important role in the fermentation of food in the GI tract of detritivorous anjak, since anaerobic fermentation is generally an important step in the digestion of plant material (van Kessel et al. 2011). Chlamydiae have been detected in diverse habitats; they are present in various animals and their major hosts, free-living amoebae, are ubiquitous (Horn 2008). Chlamydial infections of fish are emerging as an important cause of disease in new and established aquaculture industries. To date, epitheliocystis, a skin and gill disease associated with infection by these obligate intracellular pathogens, has been described in over 90 fish species, including hosts from marine and fresh water environments (Stride et al. 2014).

Vertebrate gut microbiome is a complex microbial ecosystem containing diverse and abundant bacteria, archaea, and fungi (Liu et al. 2016). The structure and composition of fish gut microbiota and their ecological function may vary with environmental locations, age, nutritional status, genotype, and morphology of the different regions of the GI tract (Ghanbari et al. 2015; Ghanbari et al. 2017; Lyons et al. 2015). To our knowledge, this is the first analysis of the gut microbial community information for the *Schizocypris altidorsalis* by high throughput sequencing approach and provide evidence of the presence of a very diverse microbial community present in this fish. The presence of many different types of bacteria in the anjak is not unexpected since it has been shown that eukaryotes with a detritivorous and planktivorous diet have a higher microbial diversity compared to omnivorous and herbivorous fish species (Ghanbari et al. 2015; Ghanbari et al. 2017; Ye et al. 2014). Many of these microbes might be of high physiological relevance for snow trout and may play key roles in the functioning of a healthy gut in these fish. Further investigations of how the bacteria coordinate in the gut microbiota and their interaction with their hosts, may offer a more in-depth insight into the complexity of this natural environment.

